# Syncmers are more sensitive than minimizers for selecting conserved *k*-mers in biological sequences

**DOI:** 10.1101/2020.09.29.319095

**Authors:** Robert C. Edgar

## Abstract

Minimizers are widely used to select subsets of fixed-length substrings (*k*-mers) from biological sequences in applications ranging from read mapping to taxonomy prediction and indexing of large datasets. The minimizer of a string of *w* consecutive *k*-mers is the *k*-mer with smallest value according to an ordering of all *k*-mers. Syncmers are a family of alternative methods which select *k*-mers by inspecting the position of the smallest-valued substring of length *s<k* within the *k*-mer. For example, a closed syncmer selected if its smallest *s*-mer is at the start of end of the *k*-mer. At least one closed syncmer must be found in every window of length (*k* – *s*) *k*-mers. Unlike a minimizer, a syncmer is identified by its sequence alone, and is therefore synchronized in the following sense: if a given *k*-mer is selected from one sequence, it will also be selected from any other sequence. Also, minimizers can be deleted by mutations in flanking sequence, which cannot happen with syncmers. Experiments on minimizers with parameters used in the minimap2 read mapper and Kraken taxonomy prediction algorithm respectively show that syncmers can simultaneously achieve both lower density and higher conservation compared to minimizers.

## Introduction

### K-mers, submers and minimizers

Next-generation sequencing (NGS) has enabled dramatic advances in fields ranging from human functional genomics (Morozova and Marra, 2008) to microbial metagenomics (Gilbert and Dupont, 2011), generating vast amounts of data currently at petabase scale and increasing exponentially (Schmidt and Hildebrandt, 2017). Computational methods for analysis of NGS data are often based on *k*-mers, short subsequences of fixed length *k*. The number of *k*-mers in an index is often very large, motivating methods for selecting subsets as a time and space optimization. I use the term *submers* for a method designed to select a common subset of *k*-mers from similar sequences. The canonical example of submers is *minimizers* (Roberts *et al*., 2004). Minimizers are used in applications including read mapping (Li, 2018), taxonomy prediction (Wood *et al*., 2019) and assembly (Sommer *et al*., 2007; Ye *et al*., 2012). Minimizers are defined by a choice of *k*, a window length *w*, and a *coding* function which assigns an integer value (*code*) to a sequence of length *k*. A coding function implies a total or partial order on the *k*-mers. The minimizer in a given window is a *k*-mer with smallest code value (ties may be handled in different ways), and the set of minimizers for a sequence is constructed by taking the union of minimizers over all windows. A simple and popular coding function is to consider letters in the nucleotide or amino acid alphabet to be base-4 or base-20 digits respectively (*lexicographic coding*). Often the coding function hashes the lexicographic code to improve the pseudo-randomness of the codes and/or to reduce the range of codes, e.g. for hash table indexes, especially with larger values of *k* where a table of size 4^*k*^ or 20^*k*^ would exceed available memory. The read mapper Winnowmap (Jain *et al*., 2020) introduced the concept of *weighted minimizers* which assigns a weight to each *k*-mer biasing its probability of being selected in the event of ties. The *density* of a submer rule is defined as the fraction of *k*-mers selected in a long random string. For given *k* and *w*, the coding function can be optimized with the goal of reducing density (Zheng *et al*., 2020; Marçais *et al*., 2018). Submers can also be defined by a *universal hitting set (UHS*) (Orenstein *et al*., 2016, 2017), an explicitly enumerated set of *k*-mers such that any sequence of length *w* must contain at least one member. Minimizers and UHS submers provide a *window guarantee*, i.e. there is necessarily at least one submer in every string of fixed length *w*. Here, I introduce syncmers, a family of alternative methods which select *k*-mers by inspecting the position of the smallest-valued substring of length *s<k* within the *k*-mer. For example, a closed syncmer selected if its smallest *s*-mer is at the start of end of the *k*-mer. At least one closed syncmer must be found in every window of length (*k* − *s*) *k*-mers (see Fig. 1 and proof in Methods).

**Figure 1.**
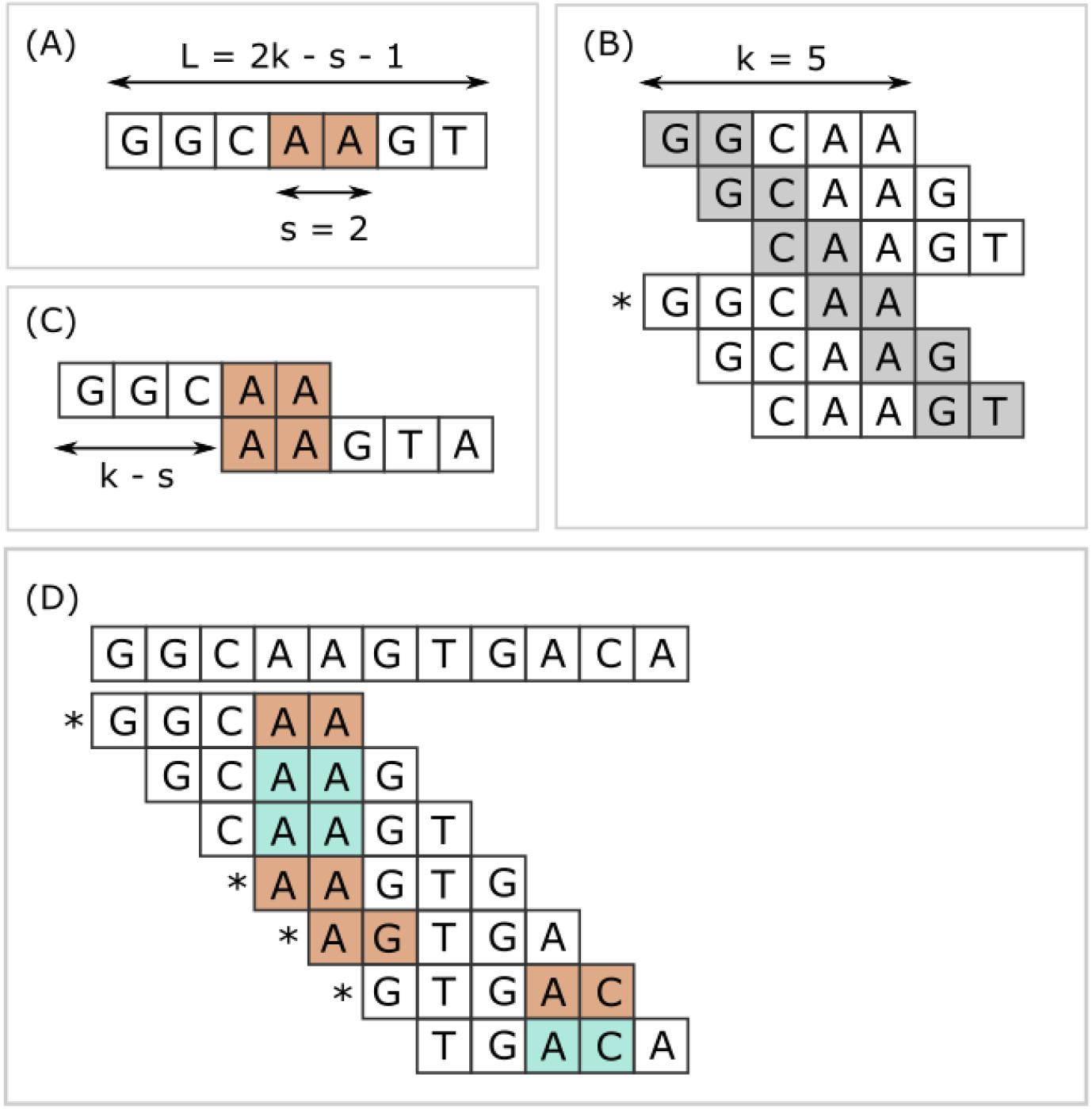
Closed syncmers. Construction of *k*=5, *s*=2 closed syncmers with lexicographic coding. A *k*-mer is a closed syncmer if its smallest *s*-mer is at the beginning or end of the *k*-mer sequence. Consider a window of three *k*-mers (length *L* = 2*k* − *s* − 1 = 7 letters) with the sequence shown in panel (A). The smallest *s*-mer is AA (orange background). Panel (B) shows the six *s*-mers in the sequence in panel (A). Each *s*-mer is shown with a gray background in the *k*-mer where it appears in the first or last position. Panel (B) illustrates that every *s*-mer in the sequence shown in (A) appears at the start or end of a *k*-mer. Therefore, regardless of which *s*-mer has the smallest value, there is a *k*-mer in the window for which this *s*-mer appears at the first or last position. In this example, AA appears at the end of GGCAA, marked with an asterisk (*), and GGCAA is therefore a syncmer. This shows that every window of length *L* must contain at least one syncmer. Note that while flanking sequence is shown in the figure, GGCAA is recognized as a syncmer from its sequence alone because its smallest 2-mer appears at the end. Closed syncmers tend to form pairs spaced at the maximum possible distance (*k* − *s*) as illustrated in panel (C). Panel (D) illustrates how *k*=5, *s*=2 closed syncmers are identified in a longer string. The smallest *s*-mer in each *k*-mer is shaded with a color. Blue background indicates that the smallest *s*-mer is not at the start or end; if it does appears at the start or end then it has an orange background and the *k*-mer is a closed syncmer (indicated by an asterisk).

### Submer conservation

Methods based on *k*-mers are generally used for comparison of similar sequences where differences may have been introduced by biological mutations and/or sequencing error. Detection of similarity by a *k*-mer requires that at least *k* letters and perhaps more are exactly conserved. Any submer is deleted if one of its letters is mutated. A minimizer is also deleted by a mutation in a position outside its *k*-mer but within its window if that mutation causes some other *k*-mer to have a smaller code. It is therefore useful to classify submers as *context-free*, i.e. selected by a rule which considers only the *k*-mer sequence, and *context-dependent*, where the rule also considers flanking sequence. Minimizers are context-dependent, while UHS submers and syncmers are context-free. Context-free submers are generally better conserved under mutation providing there are not too many overlaps between neighboring submers.

### Submer distance distribution

The length distribution of distances between consecutive submers gives insights into the advantages and disadvantages of a submer rule. An upper bound on the distance provides a window guarantee. Also desirable is that short distances have low frequencies because these correspond to submers with long overlaps which increase the rate of submer deletions under mutation. Ideally, the modal distance would have frequency 1.0 and all other distances would have zero frequency. This would uniformly tile the sequence and thereby provide a window guarantee, maximally suppress short distances, and maximize conservation. Of course, this is not achievable in practice, but provides a conceptual standard for comparison.

## Methods

### Coding function

Nucleotide *k*-mers are converted to integers by lexicographic coding. Extension to amino acid sequences is straightforward but not considered further in this report. Unless otherwise stated, lexicographic codes were hashed by murmur64 (https://en.wikipedia.org/wiki/MurmurHash).

### Minimizers

#### Definition

As noted in the introduction, variations of minimizers are possible by using different coding functions and different methods for resolving ties. In my implementation for this work, a *k*-mer is identified as a minimizer is if it has the smallest code value in any of the windows of length *k + w* - 1 letters which cover it.

#### Compression factor

Let the *compression factor* (*c*) be the number of *k*-mers divided by the number of submers selected in a long random string, so that *c* ≥ 1 and larger *c* indicates a smaller subset. This is the inverse of density as defined elsewhere in the literature (number of submers divided by number of *k*-mers). For minimizers, *c* can be estimated as follows, assuming code values are independent and identically distributed (IID) and there are no ties in a window. Consider a pair of adjacent windows in a random sequence. Sliding one position to the right, one *k*-mer (*κ*_1_) is discarded from the first window and one (*κ*_2_) is added to the second. The minimizer in the second window is different if *κ*_1_ or κ_2_ has the smallest code over both windows; otherwise the minimizer of both windows is found in their intersection and does not change. There are *w* + 1 *k*-mers in the two windows combined, and the probability that a given one of these has the smallest code is therefore *P_m_* = 1/(*w* + 1). Thus, the probability that a new minimizer is introduced by sliding the window one position is 2 *P_m_* = 2/(*w* + 1) and hence

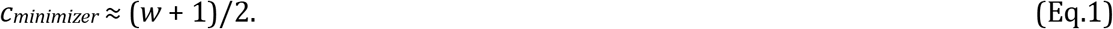

The maximum distance between consecutive minimizers, measured as the difference in their start coordinates, is *w*, as shown by the following argument. Consider a window of *w* consecutive *k*-mers where the minimizer is the first *k*-mer. Slide the window one position to the right. This must introduce a new minimizer. The maximum distance from the previous minimizer is obtained when the newly-created *k*-mer at the right-hand end of the window is the new minimizer, and this is at distance *w* from the previous minimizer which was lost at the left-hand end.

### Mincode submers

#### Definition

Mincode submers are defined by a compression factor *c* > 1 which sets a minimum value for the code, as follows: a *k*-mer *κ* is a mincode submer if code(*κ*) ≤ *H/c* where *H* is the maximum possible code. Mincode submers are context-free because no flanking sequence is considered.

#### Distance distribution

Assuming that *k*-mer codes are IID, the distance distribution for mincode submers can be calculated as follows. The probability that a given *k*-mer is selected is 1/*c*. Thus, the probability that the distance is one (immediately adjacent) is *P*(1) = 1/*c*, the probability of distance two is *P*(*r*=2) = (1 - *P*(1))*P*(1) and the probability of distance *r* is

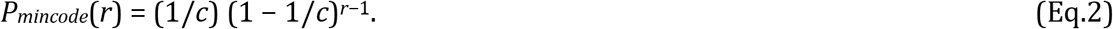

For example, with *c*=2 this gives *P*(1)=0.5, P(2)=0.25, P(3)=0.125, and with *t*=3, *P*(1)=0.33, *P*(2)=0.22, *P*(3)=0.15. For any *c* > 1, the maximum is *P*(1) and the probability of longer distances falls exponentially with *r*. With larger *c*, (1 - 1/*c*) is closer to one and the exponential falls more slowly. The maximum at *r*=1 implies that overlaps between adjacent submers are common, causing multiple submers to be disrupted by a single mutation. The slow fall of the exponential tail will cause longer gaps between covered positions. Thus, mincode submers provide no window guarantee and the distance distribution is undesirable.

### Modulo submers

#### Definition

Module submers are defined by an integer compression factor *c* > 1, as follows: a *k*-mer *κ* is a mincode submer if modulo(code(*κ*), c) = 0. Mincode submers are a particularly simple and convenient submer implementation which anecdotally I believe to be well-known in the field, though I was unable to find a reference in the literature. The properties of modulo syncmers are essentially the same as mincode syncmers with an integer value of *c*; both methods can be interpreted as selecting a randomly-chosen subset of all possible *k*-mers containing ~4^*k*^/*c* sequences.

### Closed syncmers

#### Definition

Closed syncmers are parameterized by *k*, a substring length *s < k*, and a coding function. Given a string *S*, let *μ*(*S, s*) be the *s*-mer substring with smallest code value, taking the first in left-to-right order if there is a tie. A *k*-mer *κ* is a closed syncmer if and only if *μ*(*κ, s*) is the first or last *s*-mer in *κ* (Fig. 1).

#### Window length

Consider a window *W* containing *w* = (*k* − *s*) *k*-mers. Let *σ*^+^ = *μ*(*W, s*), i.e. the first *s*-mer with smallest word value in *W*. For any *s*-mer *σ* in *W*, there is a *k*-mer *κ* such that *σ* is at the beginning or end of *κ*. Therefore *σ*^+^ is at the start or end of a *k*-mer in *W*; call this *k*-mer *κ*^+^. The *s*-mers in *κ*^+^ are a subset of *s*-mers in *W*, and therefore *σ*^+^ has the smallest word value in *κ*^+^, i.e. *μ*(*κ*^+^, *s*) = *σ*^+^ = *μ*(*W, s*). Therefore, *κ*^+^ is a syncmer, and all windows of length *w* = (*k* − *s*) *k*-mers contain at least one (*k, s*) closed syncmer. Thus, closed syncmers provide a window guarantee.

#### Compression factor

A *k*-mer has *n* = (*k* − *s* + 1) *s*-mer substrings. Under the approximation that that *s*-mer code values are IID, the probability a given *s*-mer is the smallest is 1/*n* and the probability that the first or last is the smallest is 2/*n*. Then the probability that a given *k*-mer is a closed syncmer is 2/(*k* − *s* + 1) = 2/(*w* + 1), and the compression factor for a closed syncmer is

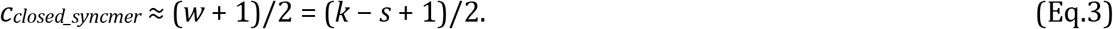

This approximation is reasonable when 4^*s*^ ≫ *k*, but breaks down when 4^*s*^ ≲ *k*. For example, if *s*=1, then a *k*-mer is a closed syncmer if the first or last letter has lowest code value (call it A), which occurs with probability 1/4 (first letter is A) + 1/4 (last letter is A) - 1/16 (subtract double-counting when both first and last letters are A) = 0.56 for any *k* > 1, and this is an under-estimate because there are additional closed syncmers containing no As. Thus, with *s*=1 the compression factor is <1/0.56 = 1.79 for any *k*.

#### Maximum distance

By the same argument as minimizers, the maximum distance between two closed syncmers is *w* = (*k* − *s*), and the minimum overlap between adjacent closed syncmers is therefore *s*.

#### Efficient spacing

There is a marked tendency for successive closed syncmers to be separated by their maximal possible distance *w* = (*k* − *s*) due to the formation of closed syncmer pairs as illustrated in Fig. 1 (C). Consider a closed syncmer *κ*_1_ for which the minimal *s*-mer *σ*_1_ is at the right-hand end, as in the upper *k*-mer shown in Fig. 1 (C), and extend the sequence with *k* − *s* additional letters to form a second *k*-mer *κ*_2_ starting with *σ*_1_. If *σ*_1_ is the smallest *s*-mer in the combined sequence *W* = *κ*_1_ ∪ *κ*_2_, then *κ*_2_ is the next closed syncmer after *κ*_1_ and the distance is (*k* − *s*). Given that *σ*_1_ has the smallest value in *κ*_1_, it is quite likely to have the smallest value in *W*, as in the example shown in Fig. 1 (C). If not, an *s*-mer *σ*_2_ with code less than *σ*1 induces a new closed syncmer before *κ*_2_ and the distance is < (*k* − *s*). Now consider a syncmer *κ*_3_ for which the minimal *s*-mer *σ*_3_ is at the start, as in the lower *k*-mer shown in Fig. 1 (C). The subsequent *κ*-mer *κ*_4_ (not shown in Fig. 1) formed by adding one more letter does not contain *σ*_1_ and must therefore have a new minimal *s*-mer *σ*_4_ which may be anywhere in *κ*_4_. If the code values of *s*-mers are assumed to be IID, then the distance distribution for this situation can be approximated by similar reasoning to the derivation of Eq.2 assuming that the probability that a given *s*-mer is the smallest in a *k*-mer is 1/(number of *s*-mers in a *k*-mer) = 1/(*k* − *s* + 1). Thus, there are two types of distance: (i) between members of the same pair, and (ii) between neighboring closed syncmers which are not paired with each other. At most half of all distances are of type (i), where the distance always (*k* − *s*). Other distances will exhibit an approximately exponential decrease given by Eq.2 with *c* = (*k* − *s* + 1) with a maximum of (*k* − *s*).

### Open syncmers

#### Definition

A *k*-mer *κ* is an open syncmer if and only if *μ*(*κ, s*) is the first *s*-mer in *κ*.

#### Compression factor

By similar reasoning to Eq.3, the compression factor is ~(*k* − *s* + 1) for open syncmers.

#### Spacing

Open syncmers provide a window guarantee, but the maximum possible distance is not easily reduced to a closed form. If a string has no open syncmer, the first occurrence of its smallest *s*-mer must be in its last (*k*-1) letters; call this the *little-endian* property. If a suffix of (*k*-1) additional letters are appended, all of the new *s*-mers must have lower values than all existing *s*-mers, otherwise the string is no longer little-endian and an open syncmer must be found. If this construction is continued, the suffixes form a strictly decreasing series which must terminate after a finite number of steps when the smallest possible *s*-mer appears. This proof shows that an upper bound on distance exists. Longer gaps are suppressed because the probability that a string is little-endian falls rapidly with *L*, which implies that the worst-case scenario suggested by the proof is highly improbable for many practically relevant values of *k* and *s*. As an example, consider the simple case *k*=3, *s*=1 with lexicographic coding. With these parameters, the longest string without an open syncmer is TTGGCCAA; it is the only 8-mer with that property and therefore occurs only once every 4^8^ = 65,536 8-mers in a random string. Thus, with open syncmers the upper bound on distance has less practical relevance than for minimizers, because with minimizers the maximum distance has the same frequency as all other distances, while with open syncmers the maximum distance is very rare.

### Circular syncmers

Circular syncmers consider a *k*-mer sequence to wrap around and thereby to contain *k* distinct *s*-mers rather than *k* − *s* + 1. Non-circular syncmers are described as *linear* if needed to distinguish these two types. For example, the 5-mer ACGTA contains 2-mers AC, CG, GT and TA if considered to be linear, plus AA if circular. With lexicographic coding, ACGTA is a (*k*=5, *s*=2) linear open syncmer because the first 2-mer AC is its smallest, but not a circular open syncmer because the smallest 2-mer (AA) is not the first. By increasing the number of *s*-mers, circularity enables higher compression for given *k* and *s*. For example, the compression factor is ~(*k* − *s* + 1) for linear open syncmers which increases to ~*k* for circular open syncmers (by similar reasoning to Eq.3). By the same construction described in the proof for linear open syncmers, circular open syncmers also provide a window guarantee, and it further follows that circular closed syncmers provide a window guarantee because they are a superset of circular open syncmers.

### Offset parameter

Syncmer variants can be defined by introducing an offset parameter *t* specifying the one-based position of the *s*-mer to be considered. For example, an open syncmer with offset *t*=2 has the smallest *s*-mer in its second position rather than the first. This has the effect of eliminating immediately consecutive (distance=1) linear syncmers, as shown by the following argument. Suppose *κ*_1_ is an open syncmer with *t*=2, and append one more letter to give a new *k*-mer *κ*_2_. The smallest *s*-mer in *κ*_1_ (*σ*^+^) must be in its second position, and this is now the first *s*-mer in *κ*_2_. Either *σ*^+^ is the smallest *s*-mer in *κ*_2_, or the new *s*-mer at the end of *κ*_2_ is the smallest; in both cases the smallest *s*-mer is not in its second position and *κ*_2_ is therefore not a syncmer. If the syncmers are circular, consecutive syncmers are strongly suppressed but not completely eliminated. Similarly, for any value of *t*, distances < *t* are suppressed. If *t* > 1, the window guarantee is lost, as shown by the counter-example of a long string of the same letter (e.g. AAAAA….) which cannot contain a syncmer. However, this could be considered a desirable property in practice by avoiding seeds in low-complexity sequence.

### Down-sampled syncmers

#### Definition

Down-sampled syncmers are defined by introducing an additional parameter *d* > 1 which enables arbitrarily large compression factors. A *k*-mer *κ* is a down-sampled syncmer if *κ* is a syncmer and *κ* is a mincode submer with factor *d*.

#### Compression factor

With a well-chosen coding function, the probability that a *k*-mer is a syncmer is effectively independent of the probability that it is a mincode submer. The probabilities are therefore multiplicative and the compression factor of a down-sampled syncmer is

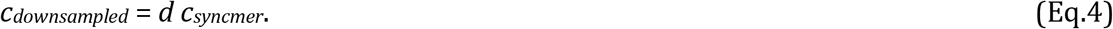

#### Spacing

Down-sampled syncmers are a subset of mincode submers with compression factor *d*, and therefore by Eq.3

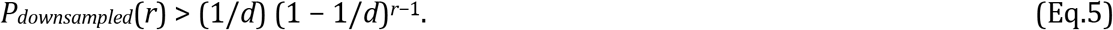

The value of *d* required to get a given compression factor *c* is smaller for down-sampled syncmers than for mincode syncmers, giving a distance distribution with faster exponential decline.

### Prefix submers

#### Definition

Let *X*(*k*) be a submer rule for *k*-mers. Define a new rule *Y*(*X, k, k*’) as follows: a *k*-mer is a *Y* submer if its first *k*’ letters (its *prefix*) is an *X*(*k*’) submer.

#### Spacing

The distance distribution of *Y*(*X, k, k*’) is identical to the distance distribution of *X*(*k*’) by construction. Informally, this is because the prefix is unchanged when letters are added, and only the prefix is considered by the rule.

#### Speed optimization of submer identification

Prefix submers enable speed optimizations. For example, suppose an application requires *k*=64 syncmers with compression *c* ≈ 3. By Eq.3, *s* ≈ *k* − 2*c* + 1 and *c* ≈ 3 can thus be achieved by (*k*=8, *s*=3) closed linear syncmers. The application can therefore use prefix closed linear syncmers with *k*=64, *k*’=8 and *s*=3 to achieve the desired compression factor. There are 65,536 possible 8-mers, which can be pre-classified and stored in a boolean vector. If nucleotide sequences are encoded with two bits per letter, the binary encoding of the 8-mer prefix can be used directly as the vector subscript because the processor can interpret the encoded 8-mer as a 16-bit integer immediately, without data type conversions. This technique reduces syncmer identification to three operations per position: a memory access, a boolean test, and a pointer increment which advances to the next position.

### Submer properties on random sequences

Submer properties including the compression factor (*c*) were measured on a random sequence *R* of length 10^6^ letters {ACGT}. The compression factor was measured as the number of *k*-mers divided by the number of submers.

### Submer conservation with random mutations

Conservation was measured by introducing random substitutions into *R* at a given frequency, giving a mutated sequence *R*’. The mutation rate is expressed as percentage identity of *R* to *R*’, e.g. frequency 0.2 is identity 80%. A submer is conserved if the *k*-mer at a given position is unchanged (no substitutions) and this *k*-mer is selected as a submer in both *R* and *R*’. The fraction of letters which are both indexed and conserved (denoted *ConsPctid*, e.g. Cons80) is defined as the number of positions appearing in conserved submers divided by the length of *R*. If a given position appears in multiple conserved submers, it is counted only once. This definition of conservation is designed to quantify the fraction of the mutated sequence covered by alignment seeds.

### Evaluation on whole-genome alignment

I implemented a pair-wise whole-genome aligner as follows. Submers and their positions in two genomes are identified and indexed in a hash table. For each pair of submers with the same sequence, one in each genome, an ungapped seed-and-extend alignment (Altschul *et al*., 1990) is constructed. The seed alignment implied by the matching submers is extended until (maximum score) - (current score) > *X*, where *X* is a fixed parameter, and the alignment with the maximum score is reported. After selecting submers, this algorithm has only three free parameters relative to the match score (1): mismatch score (−3), *X* (16) and minimum score to report an alignment (100). The minimum alignment score is set to a high value to suppress short and/or low-identity alignments where homology is ambiguous. Submer sensitivity was compared by measuring the aligned fraction (*AF*) of the input sequences obtained by different submer types. AF is calculated as (*n*_1_/*L*_1_ + *n*_2_/*L*_2_)/2 where *n*1 is the number of bases in the first genome which appear in an alignment, *L*_1_ is the length of the first genome, and similarly for the second genome. The value of AF ranges from 0 (no alignments) to 1 (all bases in both genomes appear in at least one alignment). Gapped alignments were not considered because alignment length is sensitive to gap penalty values, and the difference in AF between two submer types may therefore vary with gap penalty choice. With gapless alignments, if one submer type has substantially higher AF, it Is reasonable to conclude that this submer is more sensitive to homologous seeds. If two submer types have similar AF, then the result is inconclusive because it cannot be ruled out that one submer type would find more low-scoring alignments that are in fact homologous.

### Genome pairs

Whole-genome alignments were assessed using the bacterial assemblies shown in Table 1. Pair-wise average nucleotide identity (ANI) and aligned fraction (AF) were measured from MEGABLAST (Morgulis *et al*., 2008) alignments having *E*-values ≤ 10^-9^ and length ≥100. Gap-open and -extension penalties were set to high values to ensure that alignments were gapless. I manually selected pairs of genomes with a representative range of ANIs, preferring pairs with higher AF. These pairs were used to measure submer sensitivity in whole-genome alignment.

**Table 1.** Bacterial genome assemblies used in this study. Columns are: *Id* short identifier used in Table 3; *Assembly* NCBI assembly accession; *Description* species or strain name.

| Id | Assembly | Description |
| --- | --- | --- |
| L11 | GCA_002222595.2 | Blautia hansenii DSM 20583 |
| L13 | GCA_004295125.1 | [Clostridium] scindens ATCC 35704 |
| L14 | GCA_009684695.1 | [Clostridium] scindens |
| L15 | GCA_000178835.2 | Cellulosilyticum lentocellum DSM 5427 |
| L16 | GCA_004210255.1 | Blautia producta |
| L17 | GCA_010669205.1 | Blautia producta ATCC 27340 = DSM 2950 |
| L18 | GCA_008121495.1 | [Ruminococcus] gnavus ATCC 29149 |
| L26 | GCA_008281175.1 | [Clostridium] hylemonae DSM 15053 |
| L27 | GCA_900537995.1 | Roseburia intestinalis L1-82 |
| L30 | GCA_002234575.2 | [Clostridium] bolteae |
| L31 | GCA_000225345.1 | Roseburia hominis A2-183 |
| L41 | GCA_001689125.2 | Blautia sp. YL58 |
| L42 | GCA_001688665.2 | Lachnoclostridium sp. YL32 |
| L43 | GCA_900120345.1 | Lachnoclostridium phocaeense |
| L46 | GCA_003287895.1 | Blautia sp. N6H1-15 |
| L51 | GCA_003990395.1 | Cellulosilyticum sp. WCF-2 |
| L52 | GCA_007361935.1 | Lachnospiraceae bacterium KGMB03038 |
| F11 | GCA_003019695.1 | Fusobacterium gonidiaformans ATCC 25563 |
| F13 | GCA_000007325.1 | Fusobacterium nucleatum subsp. nucleatum ATCC 25586 |
| F14 | GCA_000158275.2 | Fusobacterium nucleatum subsp. animalis 7_1 |
| F15 | GCA_000162235.2 | Fusobacterium nucleatum subsp. vincentii 3_1_36A2 |
| F16 | GCA_000163915.2 | Fusobacterium nucleatum subsp. vincentii 3_1_27 |
| F17 | GCA_000400875.1 | Fusobacterium nucleatum subsp. animalis 4_8 |
| F18 | GCA_001296085.1 | Fusobacterium nucleatum subsp. animalis |
| F23 | GCA_001433955.1 | Fusobacterium nucleatum subsp. polymorphum |
| F24 | GCA_001457555.1 | Fusobacterium nucleatum subsp. polymorphum |
| F27 | GCA_002211645.1 | Fusobacterium nucleatum subsp. animalis |
| F35 | GCA_003019715.1 | Fusobacterium necrophorum subsp. funduliforme |
| F36 | GCA_003732505.1 | Fusobacterium necrophorum subsp. funduliforme |
| F40 | GCA_003019755.1 | Fusobacterium periodonticum |

### Representative parameters

To select parameters representative of those used in practice, I chose minimizers (*k*=15, *w*=10) and (*k*=31, *w*=15) used by minimap2 (Li, 2018) and Kraken v1 (Wood and Salzberg, 2014), respectively. Comparable syncmer parameters were identified as those giving compression factors equal or better than the minimizers with equal or better seed conservation at 90% identity (Cons90).

## Results

### Comparison of syncmers with minimap2-like minimizers

Table 2 shows properties for minimap2-like minimizers with selected *k*=15 syncmers that achieve better compression and/or better conservation. Notice for example that open syncmers with *s*=9, *t*=3 achieve compression of 7.0 vs. 5.5 for minimizers (27% lower density) with conservation of 0.312 vs. 0.301 (4% better).

**Table 2.** Syncmers with higher compression and higher conservation than minimizers with minimap2 parameters. Minimap2 uses minimizers with *k*=15, *w*=10. This table shows syncmers having both higher compression (lower density) and higher conservation than minimizers with these parameters, which are shown in the first row for comparison. Rows are sorted by increasing conservation. *OR* indicates whether the syncmers are open (*Yx*) or closed (*Nx*), and whether they are rotated (*xY*) or not (*xN*). Notice for example that open syncmers with *s*=9, *t*=3 achieve compression of 7.0 vs. 5.5 for minimizers (27% lower density) with conservation of 0.312 vs. 0.301 (4% better).

| *w* | *s* | *t* | *OR* | *c* | *Cons90* | *Cons80* |
| --- | --- | --- | --- | --- | --- | --- |
| 10 | - | - | - | 5.5 | 0.301 | 0.060 |
| - | 1 | 2 | YN | 7.0 | 0.302 | 0.062 |
| - | 1 | 2 | YY | 7.0 | 0.302 | 0.062 |
| - | 10 | - | YN | 6.0 | 0.306 | 0.064 |
| - | 10 | 5 | YN | 6.0 | 0.306 | 0.064 |
| - | 4 | - | NN | 6.0 | 0.306 | 0.063 |
| - | 1 | 2 | NY | 6.8 | 0.308 | 0.064 |
| - | 9 | 4 | YN | 7.0 | 0.308 | 0.063 |
| - | 9 | 2 | YN | 7.0 | 0.309 | 0.064 |
| - | 9 | 3 | YN | 7.0 | 0.312 | 0.064 |
| - | 10 | 4 | YN | 6.0 | 0.323 | 0.069 |
| - | 10 | 1 | YN | 6.0 | 0.324 | 0.068 |
| - | 10 | 2 | YN | 6.0 | 0.333 | 0.071 |
| - | 10 | 3 | YN | 6.0 | 0.333 | 0.071 |

### Comparison of syncmers with Kraken v1-like minimizers

Table 3 shows properties for Kraken v1-like minimizers with selected *k*=31 syncmers that achieve better compression and/or better conservation. For example, open syncmers with *s*=31, *t*=5 achieve compression of 11.0 vs. 8.5 for minimizers (29% lower density) with conservation of 0.081 vs. 0.077 (5% better).

**Table 3.**
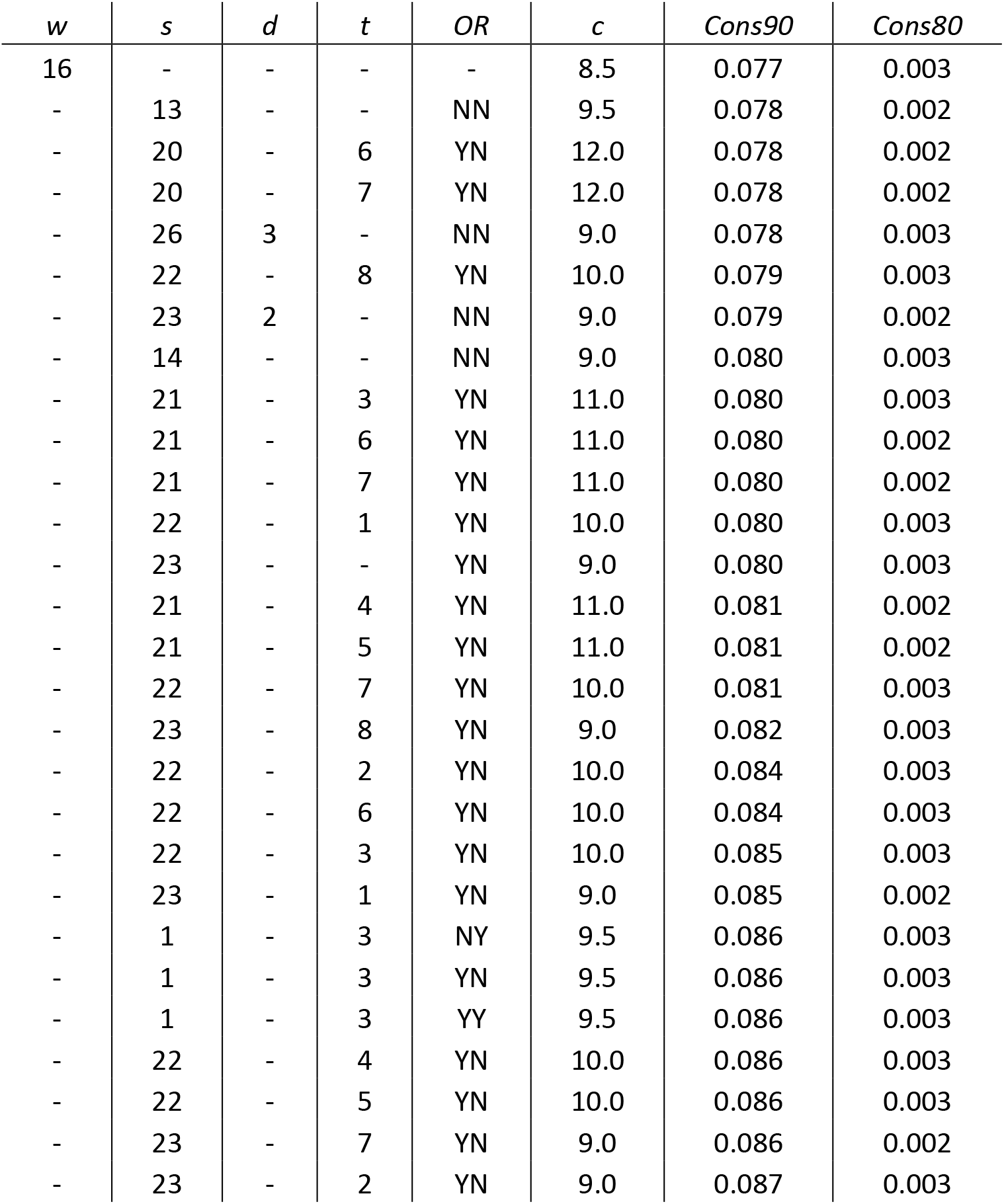
Syncmers with higher compression and higher conservation than minimizers with Kraken v1 parameters. Kraken v1 uses minimizers with *k*=31, *w*=16. For example, open syncmers with *s*=31, *t*=5 achieve compression of 11.0 vs. 8.5 for minimizers (29% lower density) with conservation of 0.081 vs. 0.077 (5% better).

| *w* | *s* | *d* | *t* | *OR* | *c* | *Cons90* | *Cons80* |
| --- | --- | --- | --- | --- | --- | --- | --- |
| 16 | - | - | - | - | 8.5 | 0.077 | 0.003 |
| - | 13 | - | - | NN | 9.5 | 0.078 | 0.002 |
| - | 20 | - | 6 | YN | 12.0 | 0.078 | 0.002 |
| - | 20 | - | 7 | YN | 12.0 | 0.078 | 0.002 |
| - | 26 | 3 | - | NN | 9.0 | 0.078 | 0.003 |
| - | 22 | - | 8 | YN | 10.0 | 0.079 | 0.003 |
| - | 23 | 2 | - | NN | 9.0 | 0.079 | 0.002 |
| - | 14 | - | - | NN | 9.0 | 0.080 | 0.003 |
| - | 21 | - | 3 | YN | 11.0 | 0.080 | 0.003 |
| - | 21 | - | 6 | YN | 11.0 | 0.080 | 0.002 |
| - | 21 | - | 7 | YN | 11.0 | 0.080 | 0.002 |
| - | 22 | - | 1 | YN | 10.0 | 0.080 | 0.003 |
| - | 23 | - | - | YN | 9.0 | 0.080 | 0.003 |
| - | 21 | - | 4 | YN | 11.0 | 0.081 | 0.002 |
| - | 21 | - | 5 | YN | 11.0 | 0.081 | 0.002 |
| - | 22 | - | 7 | YN | 10.0 | 0.081 | 0.003 |
| - | 23 | - | 8 | YN | 9.0 | 0.082 | 0.003 |
| - | 22 | - | 2 | YN | 10.0 | 0.084 | 0.003 |
| - | 22 | - | 6 | YN | 10.0 | 0.084 | 0.003 |
| - | 22 | - | 3 | YN | 10.0 | 0.085 | 0.003 |
| - | 23 | - | 1 | YN | 9.0 | 0.085 | 0.002 |
| - | 1 | - | 3 | NY | 9.5 | 0.086 | 0.003 |
| - | 1 | - | 3 | YN | 9.5 | 0.086 | 0.003 |
| - | 1 | - | 3 | YY | 9.5 | 0.086 | 0.003 |
| - | 22 | - | 4 | YN | 10.0 | 0.086 | 0.003 |
| - | 22 | - | 5 | YN | 10.0 | 0.086 | 0.003 |
| - | 23 | - | 7 | YN | 9.0 | 0.086 | 0.002 |
| - | 23 | - | 2 | YN | 9.0 | 0.087 | 0.003 |

### Comparison of distance distributions

Fig. 2 shows distance distributions for selected *k*=8 submers to illustrate how the distribution changes with submer type and parameters. As noted in the Introduction, an ideal distribution would have modal frequency 1.0, but this is not possible in practice. A desirable feature is an upper bound *w* so that all distances >*w* have frequency zero; this is equivalent to the window guarantee. Also desirable is that short distances have low frequencies because these correspond to submers with long overlaps which are more likely to be deleted under mutations. With minimizers, all distances have approximately equal frequencies (see Methods), and short distances are therefore not suppressed. Open syncmers with offset *t*>1 strongly suppress long distances and eliminate short distances, as expected (see Methods). Compare the second panel in row (A) with the second panel in row (G), which show that minimizers with *w*=8 have compression 4.5 and conservation 0.47, while open syncmers with *s*=3 and offset *t*=2 have the same conservation with substantially better compression (5.9), which can be understood as a consequence of the more desirable distance distribution of the syncmers in addition to their context-independence.

**Figure 2.**
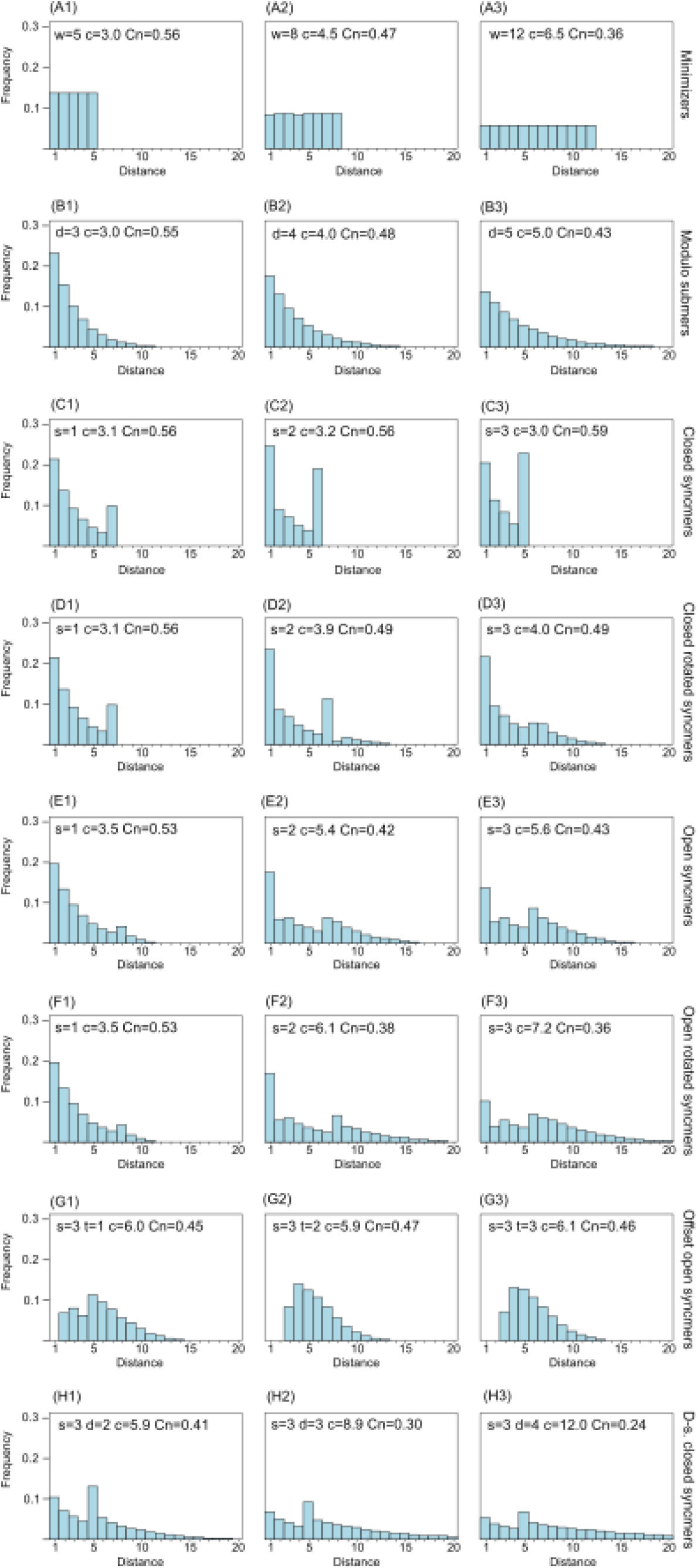
Frequency distribution of distances between consecutive submers. The histograms show spacing distributions for some representative submer types with *k*=8 including (A) minimizers, (B) modulo submers, (C) closed syncmers, (D) closed rotated syncmers, (E) open syncmers, (F) open rotated syncmers, (G) open syncmers with offset, and (H) downsampled closed syncmers. Parameters, e.g. *s* (sub-sequence length) and *t* (offset), are shown at the top of each chart, together with *c* (compression) and *Cn* (fraction of conserved letters at 90% identity) for every chart.

### Whole-genome alignment

Table 4 shows results for whole-genome alignment on 19 pairs of bacterial assemblies using (*k*=15, *w*=10) minimizers and (*k*=15, *s*=5) syncmers. On 15/19 of these pairs (79%), syncmers had higher AF (more aligned bases, indicating higher sensitivity), there was one tie, and on two pairs minimizers had 0.1% higher AF. As expected from the earlier results, syncmers show a strong tendency to have better sensitivity at lower identities. The AFs obtained with submers at low identities are much lower than MEGABLAST, indicating that many alignable segments of length ≥100 do not contain a conserved submer of either type.

**Table 4.** Results of whole-genome alignment test. ANI is average nucleotide identity, AF is aligned fraction, Min. is minimizers, Syn. is syncmers. Diff AF(%) is ((Syn. AF)*100/(Min. AF) - 100).

| Genome1 | Genome2 | Blast ANI | Min. ANI | Syn. ANI | Blast AF | Min. AF | Syn. AF | Diff. AF(%) |
| --- | --- | --- | --- | --- | --- | --- | --- | --- |
| L14 | L26 | 79.6 | 84.9 | 84.7 | 0.227 | 0.048 | 0.051 | +6.5 |
| L13 | L43 | 80.7 | 87.6 | 87.6 | 0.121 | 0.030 | 0.032 | +5.6 |
| F11 | F14 | 82.0 | 86.0 | 86.0 | 0.156 | 0.037 | 0.037 | +0.5 |
| L11 | L46 | 82.7 | 88.4 | 88.2 | 0.346 | 0.137 | 0.142 | +4.1 |
| L43 | L52 | 84.4 | 89.1 | 88.9 | 0.147 | 0.056 | 0.059 | +5.2 |
| L27 | L31 | 84.6 | 90.4 | 90.3 | 0.236 | 0.105 | 0.106 | +1.3 |
| L17 | L41 | 86.5 | 88.6 | 88.5 | 0.603 | 0.408 | 0.414 | +1.3 |
| L16 | L41 | 86.5 | 88.6 | 88.6 | 0.607 | 0.407 | 0.411 | +1.1 |
| F23 | F40 | 87.6 | 89.1 | 89.1 | 0.705 | 0.484 | 0.487 | +0.7 |
| L11 | L18 | 89.4 | 98.2 | 97.9 | 0.122 | 0.074 | 0.075 | +2.2 |
| F13 | F14 | 91.0 | 92.9 | 92.9 | 0.748 | 0.703 | 0.705 | +0.2 |
| L30 | L42 | 92.2 | 92.6 | 92.6 | 0.503 | 0.475 | 0.476 | +0.3 |
| F18 | F27 | 93.3 | 97.9 | 97.9 | 0.811 | 0.733 | 0.733 | -0.1 |
| F14 | F17 | 93.8 | 98.0 | 98.0 | 0.858 | 0.772 | 0.771 | -0.01 |
| F23 | F24 | 94.9 | 97.2 | 97.2 | 0.827 | 0.764 | 0.764 | +0.1 |
| F15 | F16 | 96.1 | 98.5 | 98.5 | 0.898 | 0.860 | 0.859 | -0.1 |
| L15 | L51 | 96.7 | 98.3 | 98.3 | 0.826 | 0.800 | 0.800 | +0.1 |
| F35 | F36 | 98.4 | 99.1 | 99.1 | 0.869 | 0.832 | 0.832 | 0.0 |
| L13 | L14 | 99.4 | 99.7 | 99.7 | 0.786 | 0.775 | 0.778 | +0.4 |

### Maximum distance under mutation

Table 5 shows maximum distances for minimap2-like and Kraken-like minimizers compared to selected syncmers tested on random sequences of length 1 M (comparable to a short bacterial genome) and 100 M (short mammalian chromosome) at identities of 95% and 90%. For *k*=15, these are linear open syncmers with *s*=9 and offset *t*=4; for *k*=31 they are linear open syncmers with *s*=21, *t*=5. These were chosen because they have both better compression and better conservation than minimizers (Tables 2 and 3). Neither of these syncmer types has a window guarantee. In identical sequences, the minimizers have a maximum distance (window size *w*) of 10 (*k*=15) and 16 (*k*=31), but under mutation the maximum distance is much greater, ranging from ~20*w* (*k*=15, 1Mb, 95%id) to ~300*w* (*k*=31, 100Mb, 90%id). Thus, the window guarantee is degraded by more than an order of magnitude even at a relatively modest 5% mutation rate in a relatively short sequence (1Mb), getting worse as the identity is reduced and/or the sequence length is increased. As shown in Table 5, syncmers often have lower maximum distance under mutation despite the lack of a window guarantee, which can be understood as a consequence of their better conservation. This observation calls into question the practical relevance of the window guarantee (see Discussion).

**Table 5.**
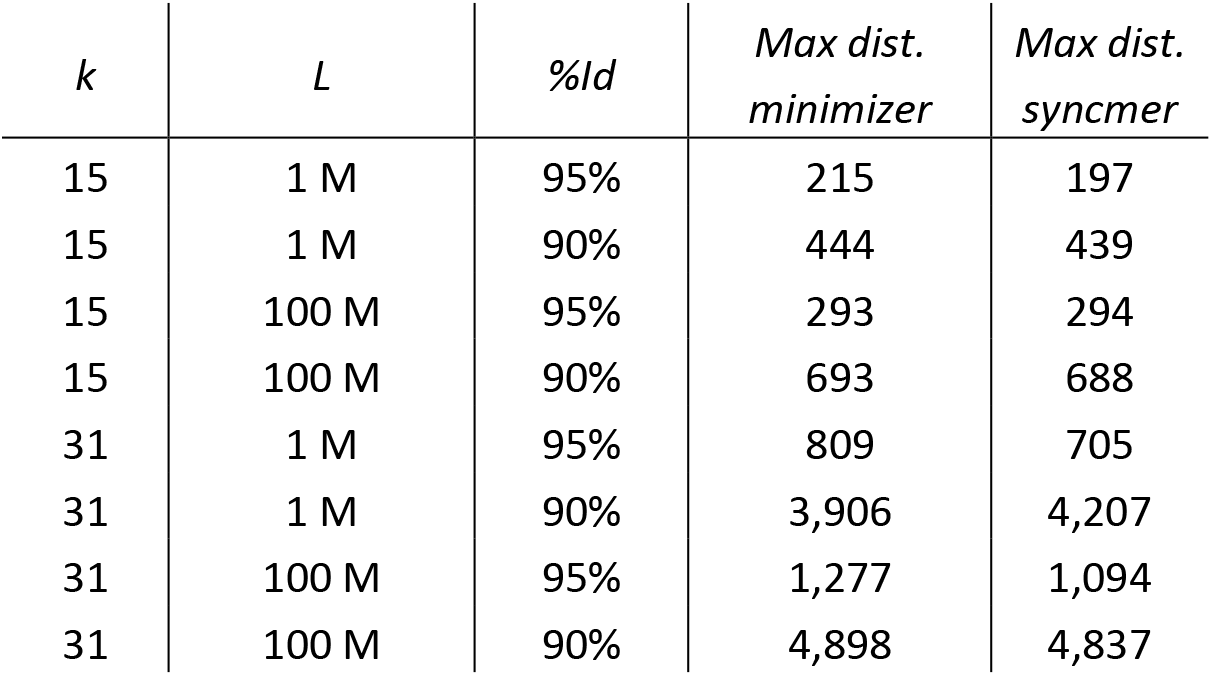
Maximum distances under mutations. Maximum distance for minimap2-like and Kraken-like minimizers compared to selected syncmers on random sequences of length (*L*) 1 M and 100 M at identities of 95% and 90%. For *k*=15, these are open syncmers with *s*=9 and offset *t=4;* for *k*=31 they are open syncmers with *s*=21, *t*=5. Neither of these syncmer types has a window guarantee. In identical sequences, the minimizers have a window size of 10 (*k*=15) and 16 (*k*=31), but under mutation the maximum spacing is much greater, even at 95% identity. Syncmers often have lower maximum spacing under mutation despite the lack of a window guarantee.

| *k* | *L* | %*Id* | *Max dist.*<br>*minimizer* | *Max dist.*<br>*syncmer* |
| --- | --- | --- | --- | --- |
| 15 | 1 M | 95% | 215 | 197 |
| 15 | 1 M | 90% | 444 | 439 |
| 15 | 100 M | 95% | 293 | 294 |
| 15 | 100 M | 90% | 693 | 688 |
| 31 | 1 M | 95% | 809 | 705 |
| 31 | 1 M | 90% | 3,906 | 4,207 |
| 31 | 100 M | 95% | 1,277 | 1,094 |
| 31 | 100 M | 90% | 4,898 | 4,837 |

### Parameter sweep

Supplementary Table 1 gives quality metrics for a wide range of submers, enabling programmers and end-users to choose parameters giving a preferred trade-off between compression factor and conservation.

## Discussion

### Connection to the miniception algorithm

The miniception algorithm (Zheng *et al*., 2020) introduced the concept of a *charged context*, a window of length (*w*_0_+1) *k*_0_-mers such that the (*k*_0_, *w*_0_) minimizer in the first window is different from the minimizer in the second window. This can happen only if the first *k*_0_-mer in the first window is the smallest or if the last *k*_0_-mer in the second window is the smallest over the combined window, which in turn implies that one of the two *w*_0_-mers is a (*k=w*_0_, *s=k*_0_) closed linear syncmer in my terminology. However, (Zheng *et al*., 2020) did not suggest selecting charged contexts as submers; their proposal was to construct charged contexts as an intermediate step to selecting minimizers with a lower density than previous methods.

### Submer rules

The modulo submer method seems obvious and has probably been reinvented many times. It is a natural standard of comparison for more elaborate methods such as minimizers, and it is puzzling that no previous comparison has been published to the best of my knowledge. Possibly the fact that arbitrarily long gaps can occur, albeit with exponentially low probability, was considered a self-evident disadvantage, and naively this appears difficult to overcome without considering flanking sequence in a window. The syncmer method demonstrates that correlations between neighboring *k*-mers can be exploited to achieve a window guarantee with a context-free rule. Many variations on this technique are possible, allowing fine-tuning of characteristics such as the spacing distribution at a desired density. Perhaps surprisingly, it is possible to obtain a window guarantee even for submers that are sparse in the sense that that they often do not overlap; this can be understood as additionally exploiting the finite (and typically small) number of distinct *s*-mers (see the “strictly decreasing series” proof for open syncmers). Introducing an offset gives up the window guarantee in favor of suppressing short distances (i.e., long overlaps) between neighboring syncmers, which improves conservation. The improved conservation achieved by this technique tends to compensate for its lack of a strict window guarantee so that it may exhibit a shorter maximum distance under mutation than similar rules which do have a guarantee for identical sequences.

### Context-free submers do not have edge bias

A *k*-mer at the start or end (edge) of a sequence appears in only one window, and thus has a probability 1/*w* of being a minimizer which is roughly half the probability 2/(*w* + 1) for a *k*-mer far enough from the edge that it appears in *w* windows (see Eq.1). This issue can be significant in practice for short sequences such as next-generation reads, e.g. for identifying overlaps in assembly. This was noted in (Roberts *et al*., 2004), where it was proposed that the problem can be addressed by introducing a special case at sequence terminals they called *end-minimizers*. This issue does not arise with context-free submers such as syncmers.

### Choosing submer parameters

An application must choose submer parameters such as *k* and *w* or *s*. Typically, these choices are driven by maximizing sensitivity and reducing computational resources, which are conflicting goals. The primary parameter is *k*, which gives a trade-off between sensitivity and specificity. Small *k* is more sensitive because mutations are less likely to delete a *k*-mer, but gives more seeds which increases processing time and may result in more false positive predictions. Larger *k* improves specificity (seeds are more likely to be homologous) and saves time (fewer seeds will be found in diverged sequences), but reduces sensitivity because more *k*-mers are deleted by mutations. As an example, bacterial genomes are ~10Mb when both strands are included. If *k*≤11, then the probability of finding any given *k*-mer in a bacterial genome is ~1 because 4^11^ ≈ 4Mb. Therefore, an application such as Kraken which requires submers to be unique within a clade must set *k* ≫ 11 to achieve acceptable specificity. By contrast, a bacterial whole-genome aligner might reasonably set *k* ~ 11 because the number of false positive seeds between one pair of genomes should be tolerably low. In the following, I will assume that *k* has been fixed based on such considerations and focus on the choice of submer rule and the rule’s parameters.

### The window guarantee is a weak heuristic

The window guarantee is designed to ensure that similar subsequences have seeds, i.e. conserved submers. This is a reasonable heuristic, but when mutations are introduced, the guarantee is necessarily lost for any type of submer. As illustrated by the examples in Fig. 3, submers, especially context-dependent submers such as minimizers, are deleted much more rapidly than letters as sequences mutate, with a large majority deleted at identities around 90% or less. Therefore, even if a submer rule has a window guarantee for identical sequences, its seeds will often have gaps ≫ *w* when sequences have ≲95% identity (Table 5). If the primary goal of the submer method is to identify seeds in homologous sequences, then the window guarantee does not provide an upper bound on spacing; it is at best a weak heuristic and should not be regarded as an essential property. If a rule without a window guarantee conserves more seeds at a given density, then it is likely to give better sensitivity in many practical applications.

**Figure 3.**
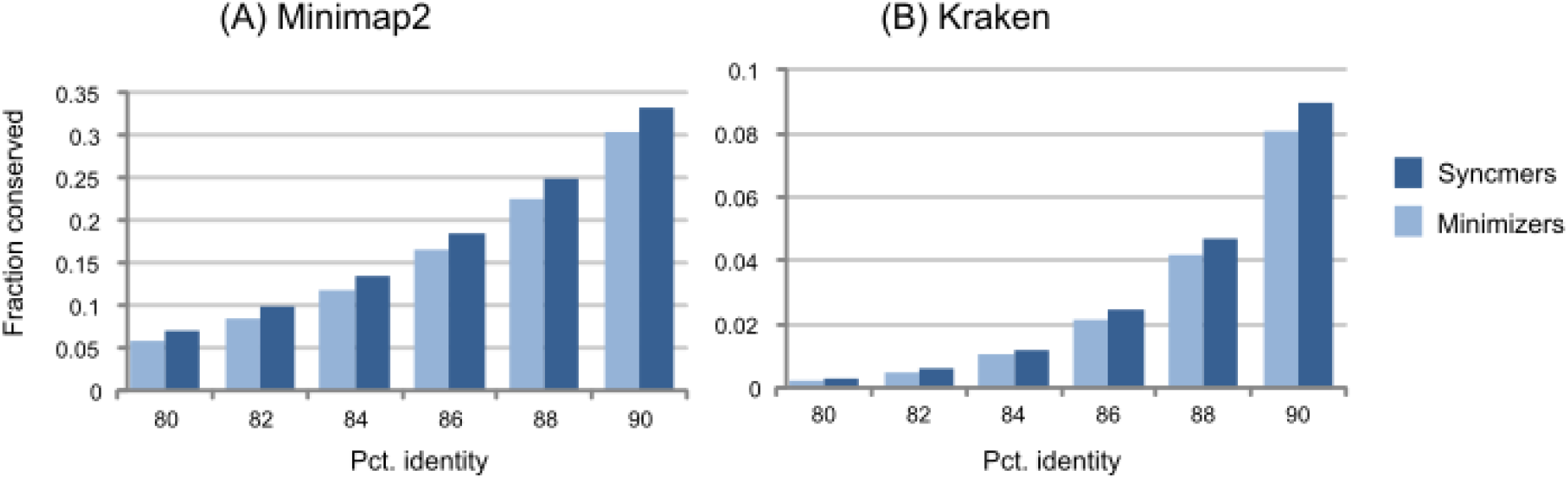
Conservation as a function of identity. The histograms show fraction of conserved letters at identities from 80 to 90% for closed syncmers and minimizers comparable to those used in minimap2 (panel A) and Kraken (panel B). Notice that the large majority of submers are deleted at these identities.

### Density is not the appropriate optimization metric

Several recent papers (see Zheng *et al*. 2020 and references therein) have focused on minimizing the density of minimizers for given *k* and *w*. This would be an appropriate optimization strategy if submers were used to find identical longer substrings in different sequences, but this is rarely the primary goal of an application and other methods are better suited to this task (e.g., Burrows-Wheeler indexes). In practice, most algorithms use minimizers to find homologous sequences which may have diverged due to mutations and/or sequencing error; for example, minimap2 and Kraken are algorithms of this type. Such applications generally aim to maximize sensitivity to diverged sequences under time and space constraints. If two minimizer rules with the same *k* and *w* have different density, then the rule with higher density will conserve more submers and is therefore more sensitive. Tuning minimizer density is thus similar to tuning the window length in that it gives a trade-off between sensitivity and index size. If lower minimizer density is needed to save memory or increase speed, then the window length could be increased, or the rule could be changed to one giving lower density. Either strategy could give better conservation, depending on the rules. For most applications, the appropriate optimization metric for submers is conservation, which should be maximized while keeping density fixed and allowing other parameters (e.g, *w*) to vary. Density should be held fixed because this parameter determines time and space resources for a given *k*. Maximizing conservation at a fixed density will tend to maximize the sensitivity achieved by an application with resources determined by the density. Conservation should generally be assessed by the number of conserved letters covered by a seed rather than the number of conserved submers to avoid double-counting when submers overlap.

### Well-conserved submers

If it is accepted that the optimization goal is maximizing conservation for fixed *k* and density, then context-free submers have a natural advantage over minimizers because a mutation anywhere in the window can delete its minimizer, while context-free submers are deleted only when a mutation occurs within the *k*-mer. The advantage is greater at lower identities, because mutations in minimizer flanking sequences are more common, and also at lower densities (for a given minimizer rule) because lower densities require longer windows, which result in more deleted minimizers due to flanking mutations. The apparent advantage of the window guarantee is therefore a liability in practice for minimizers. If the desired density is high enough that it can be achieved by closed syncmers, then they offer an unambiguously superior alternative to minimizers for a wide range of densities and *k* values because they simultaneously achieve lower density and higher conservation (Tables 2 and 3 and Supplementary Table 1) while giving a similar window guarantee. Strictly, these results are established only for my own implementation of minimizers, but they almost certainly apply to minimizer variants with different coding functions or tie-breaking rules if submers with the same density are compared rather than the same window length. At lower densities, variants such as open syncmers with an offset can again simultaneously achieve lower densities and higher conservation compared to minimizers (Tables 2 and 3 and Supplementary Table 1). Some variants lack a window guarantee, but these syncmers are nevertheless clearly superior to minimizers unless a practically relevant advantage of the guarantee can be shown. If the roles were reversed and syncmers had been proposed 16 years before minimizers, the window guarantee might now be perceived not as self-evidently advantageous, but rather as a theoretical abstraction valid only for identical sequences which is insufficient to justify accepting lower sensitivity in practice.

## Conclusions

Syncmers are a family of novel methods for selecting *k*-mers which are shown to have several properties that will be useful in practical applications. Like minimizers, they can provide a guarantee that at least one syncmer will be selected in a window of a predetermined length. Unlike minimizers, syncmers are robust against mutations in flanking sequence and are better conserved in mutated sequences.

